# The Role of Orexin-1 Receptor Signaling in Demand for the Opioid Fentanyl

**DOI:** 10.1101/533059

**Authors:** Jennifer E. Fragale, Caroline B. Pantazis, Morgan H. James, Gary Aston-Jones

## Abstract

The orexin system is a potential treatment target for drug addiction. Orexin-1 receptor (OxR1) antagonism reduces demand for cocaine and remifentanil indicating that orexin-based therapies may reduce demand for many classes of abused drugs. However, pharmacokinetics vary greatly among opioids and it is unclear if OxR1 antagonism would reduce demand for all opioids, particularly ones with particularly high abuse liability. Here, we establish a behavioral economics (BE) procedure to assess the effects of OxR1 antagonism on demand for the highly abused opioid fentanyl. We also investigate the utility of our novel procedure in predicting OxR1 antagonism efficacy and relapse propensity. The OxR1 antagonist SB-334867 (SB) increased demand elasticity (α; decreased motivation) for fentanyl without affecting preferred consumption of fentanyl at null cost (Q_o_). Baseline α values predicted SB efficacy, such that SB was most effective at increasing demand elasticity in highly motivated rats. SB also attenuated cue-induced fentanyl seeking and baseline α values predicted the amount of reinstatement behavior. These results highlight the promise of the orexin system as a treatment target for opioid addiction and emphasize the usefulness of BE procedures in the study of opioid abuse.

## Introduction

Opioid addiction is a chronic and relapsing disorder that has become a national epidemic. Six out of every 10 overdose deaths in the United States involve opioids (Rudd et al., 2016). Among the opioids most frequently abused by addicts is fentanyl, a short acting synthetic opioid that is 50 times stronger than heroin (Gladden et al., 2016; Peterson et al., 2016). The rapid increase in opioid abuse emphasizes the need for novel approaches to study addiction and a greater understanding of the neural mechanisms underlying this disorder.

Pathological demand for drug taking is a key characteristic of addiction that predicts symptom severity and treatment outcomes (Bruner and Johnson, 2014; MacKillop et al., 2018; MacKillop et al., 2008; Pickover et al., 2016). Behavioral economics (BE) procedures provide a quantitative analysis of drug demand that is highly translational (Bickel et al., 1993; Bruner and Johnson, 2014; MacKillop et al., 2018; MacKillop et al., 2008; Murphy et al., 2009; Pickover et al., 2016). Drug demand can be assessed in humans using hypothetical purchasing tasks (MacKillop et al., 2008; Murphy et al., 2009) and in preclinical models using operant tasks (Bentzley et al., 2013; Bentzley et al., 2014; Porter-Stransky et al., 2017). BE procedures quantify demand by plotting drug consumption as a function of price. A demand curve can then be fitted to the data using an exponential demand equation (Hursh and Silberberg, 2008). This equation yields two important variables. Qo describes the theoretical consumption of a reinforcer when no effort is required and is also used as a measure of hedonic set point. Alpha (α) is a measure of demand elasticity and can be used as an inverse measure of motivation. Preclinical within-session BE procedures are readily used to assess pathological demand for a variety of positive and negative reinforcers (Bickel et al., 1993; Bickel et al., 2010; Christensen et al., 2008; Fragale et al., 2017; Hursh, 1991). However, few preclinical studies have applied BE procedures to the study of opioid addiction.

The orexin (hypocretin) system is a promising treatment target for drug addiction (Baimel et al., 2015; James et al., 2017; Kenny, 2011; Lawrence et al., 2006; Sharf et al., 2010b) and orexin-based therapies have been listed by the National Institute on Drug Abuse’s (NIDA) Division of Therapeutics and Medical Consequences as a high priority treatment target for opioid abuse (Rasmussen et al., 2018). Orexins are a pair of neuropeptides exclusively produced in the hypothalamus (de Lecea et al., 1998; Sakurai et al., 1998). Orexins regulate a wide range of behaviors by directly acting on two G-coupled protein receptors, orexin-1 (OxR1) and orexin-2 (OxR2) receptors (Sakurai et al., 1998). OxR1 signaling has been implicated in a variety of drug seeking behaviors and plays a clear role in opioid addiction (Harris et al., 2007; Narita et al., 2006; Sharf et al., 2010a). Initial studies found that stimulation of a lateral subpopulation of orexin-producing neurons reinstated extinguished morphine place preference and that this effect was blocked by systemic pretreatment with the selective OxR1 antagonist SB-334867 (SB; (Harris et al., 2005). Systemic pretreatment with SB also reduced heroin self-administration and attenuated cue-elicited reinstatement of heroin seeking (Smith and Aston-Jones, 2012). More recently, we used the within-session BE procedure to show that OxR1 blockade decreases demand for the short acting opioid remifentanil (Porter-Stransky et al., 2017). Together with evidence that OxR1 antagonists are also effective at blocking addiction-related behaviors across other drugs of abuse tested including cocaine, alcohol, and nicotine, there is significant speculation that orexin-based therapies might represent a novel treatment approach for addiction in general, regardless of drug class (James et al., 2017; Kenny, 2011; Lawrence et al., 2006).

Despite this, the effects of Or1R blockade on fentanyl demand is yet to be tested. This is important because the pharmacokinetics are different among opioids (Bostrom et al., 2006; Inturrisi, 2002; Volpe et al., 2011), and because fentanyl is a major contributor to the current opioid epidemic (Gladden et al., 2016; Peterson et al., 2016). Here, we establish a novel within-session BE procedure to study the usefulness of OxR1 antagonism in reducing demand for fentanyl. We also assess the effectiveness of our novel within-session BE procedure in predicting OxR1 antagonism efficacy as well as relapse propensity.

## Methods

### Animals

Male Sprague Dawley rats were obtained from Charles River Laboratories (Kingston, NY) at 250-300g. Animals were pair-housed under a 12:12 h reverse light cycle (lights off 8:00 and on at 20:00). Animals were given *ad libitum* access to food and water. All procedures were approved by Rutgers University New Brunswick Institutional Animal Care and Use Committee and conducted in accordance with their guidelines.

### Surgery

Animals were anesthetized with 2% isoflurane, administered the analgesic rimadyl (5 mg/kg, s.c.), and implanted with chronic indwelling catheters jugular vein catheters as previously described (Smith and Aston-Jones, 2012). Following surgery, rats were flushed daily with the antibiotic cefazolin (0.1 mL; 100mg/mL) and heparin (0.1 ml; 100 U/ml). Rats were allowed one week to recover prior to self-administration training.

### Drugs

Fentanyl HCl powder and the selective OxR1 antagonist SB-334867 were generously donated by the National Institute of Drug Abuse. Fentanyl HCl powder was dissolved in 0.9% sterile saline to a concentration of 8μg/ml. SB-334867 was prepared as previously described and injected i.p. at a volume of 4.0 ml/kg (Bentzley and Aston-Jones, 2015; Porter-Stransky et al., 2017; Smith and Aston-Jones, 2012).

### Self-Administration Training

Rats were trained in Med Associates operant boxes (Med Associates, St Albans, VT, USA) fitted with 2 levers (active and inactive), a stimulus light located directly above the active lever, speaker, and house light. Operant boxes were housed in individual sound-attenuating chambers, and all boxes were controlled by Med-PC IV software (Med Associates, St Albans, VT, USA). Rats were initially trained in 2 hr sessions on a fixed-ratio 1 (FR-1) schedule. A response on the active lever resulted in a 3.6s infusion of 0.5μg of fentanyl paired with stimulus light (white light) and tone (78 dB, 2900 Hz). Each infusion was followed by a 20-sec time out signaled by termination of the house light. Responses on the inactive lever were not reinforced (no infusion or cues). Rats were trained for a minimum of 10 sessions and to a criterion of > 25 infusion for 3 consecutive sessions.

### Within-Session Behavioral Economics Procedure

Demand for fentanyl was determined using a within-session behavioral economics procedure consistent with previous studies (Bentzley and Aston-Jones, 2015; Bentzley et al., 2014; Porter-Stransky et al., 2017). Briefly, animals were tested in 110-minute sessions where the dose of fentanyl per infusion was decreased in successive 10-minute bins on a quarter logarithmic scale (1.12, 0.63, 0.36, 0.20, 0.11, 0.06, 0.04, 0.02, 0.01, 0.006, 0.004 μg fentanyl per infusion). Demand curves were generated after each session for individual subjects by fitting data to an exponential demand equation (Hursh and Silberberg, 2008). Consistent with our BE procedure for remifentanil (Porter-Stransky et al., 2017), curve fitting was performed by including all data points up until two bins past the point of maximal responding (P_max_).

Rats were trained for a minimum of 6 days and until stable demand was observed before any pharmacological manipulation. Animals were considered stable when demand parameters(Q_o_ and α values) differed by less than 25% across three consecutive sessions. Once stable, rats were given an i.p. injection of vehicle or SB (3, 10, and 30 mg/kg) 30min prior to testing. Rats were given a minimum of 3 days between testing to insure a return to baseline demand.

Treatment order was assigned pseudorandomly with all rats receiving 30 mg/kg on their final test. This was done to alleviate any concerns regarding a possible long duration of effects with 30 mg/kg SB (Brodnik et al., 2018).

### Fentanyl Demand Locomotor Control

Following BE testing, a subset of rats were trained to nose poke for sucrose (45 mg sucrose pellets, Test Diet, Richmond, IN, USA) on an FR-1 schedule. During each session, rats also received their preferred consumption of fentanyl (i.v.) as determined by their baseline Qo. This was achieved by altering infusion duration and frequency to mimic responding during individual Pmax bins (the final bin that preferred consumption is maintained). Rats were trained for a minimum of 5 sessions and until number of responses differed by less than 25% across three consecutive sessions. Vehicle or SB (30 mg/kg; the highest dose tested in demand experiments) was given in a within-subjects, counterbalanced design with a minimum of 3 days between tests.

### Extinction and Cue-Induced Reinstatement

After BE testing, a subset of rats was exposed to 2-hr extinction sessions during which responses on the active lever were no longer reinforced (no infusion or cues). Rats were trained for a minimum of 7 sessions, and until responding in the last 3 sessions was ≤ 25 presses. During reinstatement tests, presses on the active lever were paired with fentanyl-associated cues (light and tone). Pretreatment with vehicle or SB (10 and 30 mg/kg) was given in a within-subjects, counterbalanced design with a minimum of 3 days between reinstatement tests.

### Cued-Reinstatement Locomotor Control

After their final reinstatement test, the same rats were trained to lever press for sucrose (45 mg sucrose pellets, Test Diet, Richmond, IN, USA) on a FR-1 schedule during 2-hr sessions. Presses on the active lever resulted in pellet delivery that was not paired with cues. Each pellet delivery was followed by a 20-sec time out signaled by termination of the house light. Presses on the inactive lever were not reinforced. Rats were trained for a minimum of 5 sessions and until stable responding was observed. Rats were considered stable when the number of pellets obtained differed by less than 25% across three consecutive sessions. Vehicle or SB (30 mg/kg; the highest dose tested) was given in a within-subjects, counterbalanced design with a minimum of 3 days between tests.

### Data Analysis

Data are expressed as mean values ± 1 standard error of the mean. Statistics were performed using GraphPad Prism for Mac (Version 7, GraphPad Software Inc., La Jolla, CA) with an α level of 0.05. Violations of sphericity determined by Mauchly’s Test were adjusted using the Greenhouse-Geisser correction. FR-1 self-administration and extinction data were analyzed using mixed-design ANOVAs with Bonferroni corrections as *post hoc* tests. One-way repeated measures ANOVAs were used to examine the effect of SB on fentanyl demand and reinstatement. Dunnett's multiple comparisons test were used as *post hoc* tests when permitted. Data from locomotor controls were analyzed using a paired samples t-tests. Two-tailed Pearson correlations were used to investigate the relationship between baseline demand and SB efficacy as well as the relationship between baseline demand and reinstatement.

## Results

### Economic demand profile of fentanyl

Rats (n=18) were first trained to self-administer fentanyl on a FR-1 schedule in 2 hr sessions. Figure 1A shows self-administration data from the last 10 acquisition sessions prior to behavioral economics training. The means (± SEM) for the final day of self-administration training were 39.7 (±4.5) infusions, 64.0 (±9.4) active lever presses and 3.3 (±0.9) inactive lever presses. Rats readily learned to discriminate between the active and inactive lever (main effect of Lever Type F_1, 17_=18.75, p=0.0005 and a Lever Type x Session interaction F_9, 153_=2.326) with rats responding more on the active lever across the final 9 sessions (Bonferroni correction, p<0.05).

**Figure 1.**
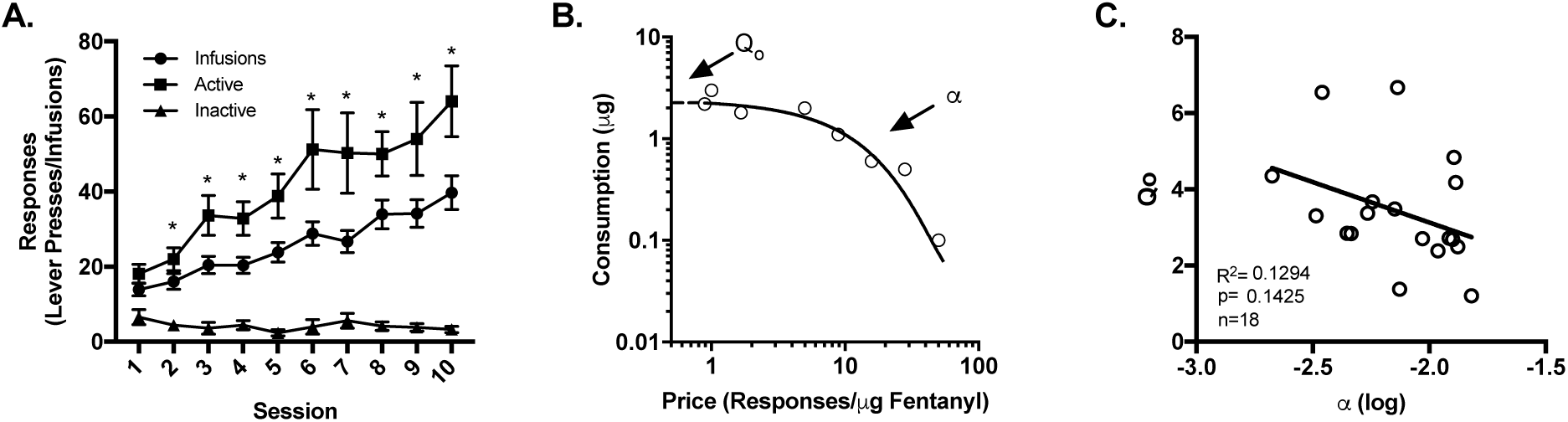
Fentanyl self-administration and demand. A) Rats were first trained to self-administer fentanyl on an FR-1 schedule. Rats readily learned to discriminate between the active and inactive levers making more active vs inactive lever pressed across the final 9 self-administration sessions. B) Shown is a demand curve generated from a representative subject from a single session. C) α and Qo are independent parameters as demand elasticity (α) was not significantly correlated with consumption at null cost (Q_o_). * p<0.05.

Once fentanyl self-administration was acquired, rats (n=18) were trained on a within-session behavioral economics (BE) procedure for fentanyl. Figure 1B shows a demand curve generated with this procedure for a representative rat and session. Overall, curve fits were very good, with r^2^ values for demand curves between 0.72 and 0.99. Mean r^2^ values were 0.91 (± 0.008). The mean (± SEM) baseline α and Qo values were 0.0834 (± 0.0009) and 3.433 (±0.351), respectively. Consistent with demand for cocaine, no relationship was observed between demand elasticity (α) and preferred intake at null cost (Q_o_; Figure 1C; R^2^ = 0.1294, p=0.129).

### SB decreases demand elasticity for fentanyl

To determine if signaling at OxR1s regulate demand for fentanyl, rats (n=14) were treated with a systemic injection of vehicle or SB (3, 10, or 30 mg/kg) prior to BE testing. Figure 2A shows representative demand curves from a subject treated with vehicle and SB (30 mg/kg) prior to BE testing. SB increased demand elasticity (increased α; decreased motivation; Figure 2B; F_2.66, 34.53_ = 8.187, p=0.0007). *Post hoc* analysis revealed that pretreatment with SB (30 mg/kg) significantly increased demand elasticity to vehicle pretreatment. (Dunnett's multiple comparisons test, p=0.005). In contrast, systemic SB pretreatment had no effect on fentanyl consumption at null cost (Qo; Figure 2C; F_2.95, 38.33_ = 1.155, p=0.3391).

**Figure 2.**
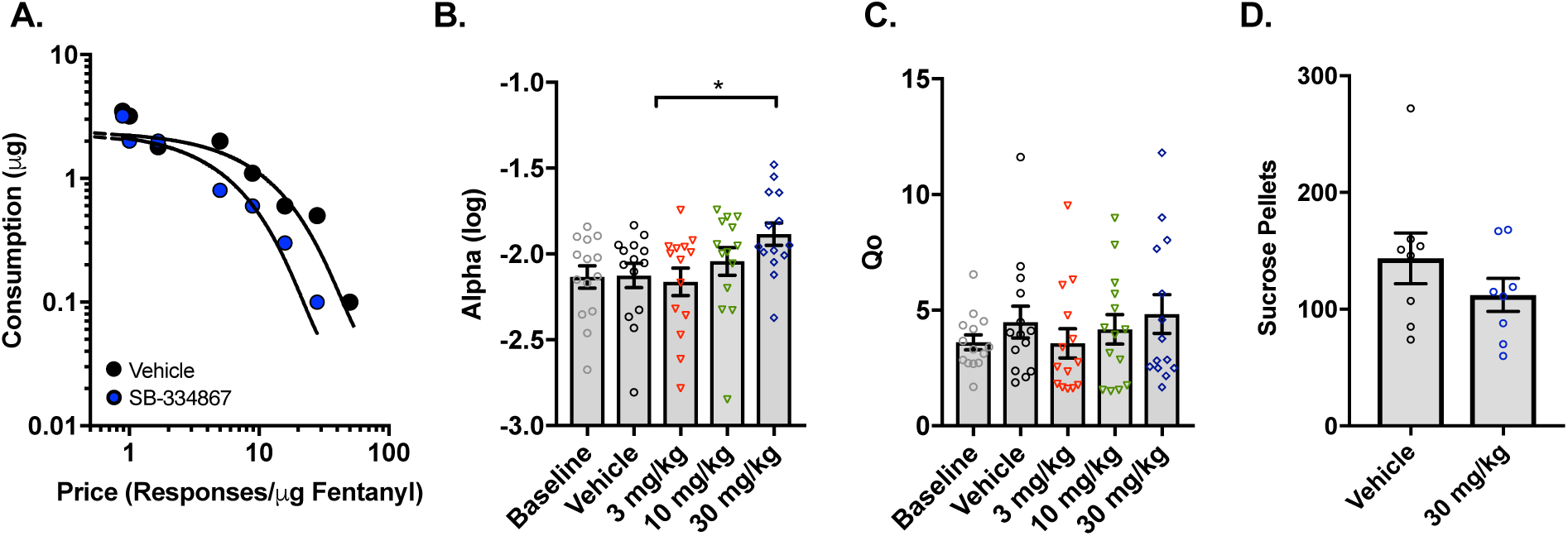
OxR1 antagonism decreases demand for fentanyl. Rats were given vehicle or SB (3, 10, or 30 mg/kg) prior to BE testing. A) Show are representative demand curves from a single subject that received vehicle and SB (30mg/kg) B) SB increased demand elasticity (increased α), but was most effective at 30 mg/kg. C) SB had no effect on preferred consumption of fentanyl at null cost (Qo). D) To determine if SB impaired locomotor activity, a subset of rats was trained to nosepoke for sucrose on a FR-1 schedule while receiving their preferred consumption of fentanyl (i.v.). SB (30 mg/kg) had no significant effect on sucrose consumption compared to vehicle treated rats. * p<0.05

Following BE testing, a subset of rats (n=8) were trained to nose poke for sucrose on an FR-1 schedule while receiving their preferred consumption of fentanyl (i.v.) as determined by their baseline Qo. This was done to determine if the effects of SB on demand elasticity were due to sedation or motor impairment. SB (30 mg/kg) did not alter nose poking for sucrose in rats receiving their preferred fentanyl doses (Figure 2D; t_7_=1.958, p=0.0911).

### Baseline α values predict SB efficacy

We previously found that baseline demand elasticity for cocaine predicts the extent to which SB reduces motivation for cocaine, such that SB is most effective in rats with inelastic demand (low α) (James et al., 2018a). Accordingly, here we examined the relationship between baseline demand elasticity for fentanyl and the extent to which SB affected motivation during BE testing. Baseline α was negatively correlated with SB efficacy (30mg/kg) such that SB was most effective at increasing demand elasticity in highly motivated rats (low α; Figure 3A; R^2^= 0.3462, p=0.0269) No relationship was observed between baseline Qo values and the efficacy of SB in increasing demand elasticity (Figure 3B; R^2^=0.00052, p=0.8047).

**Figure 3.**
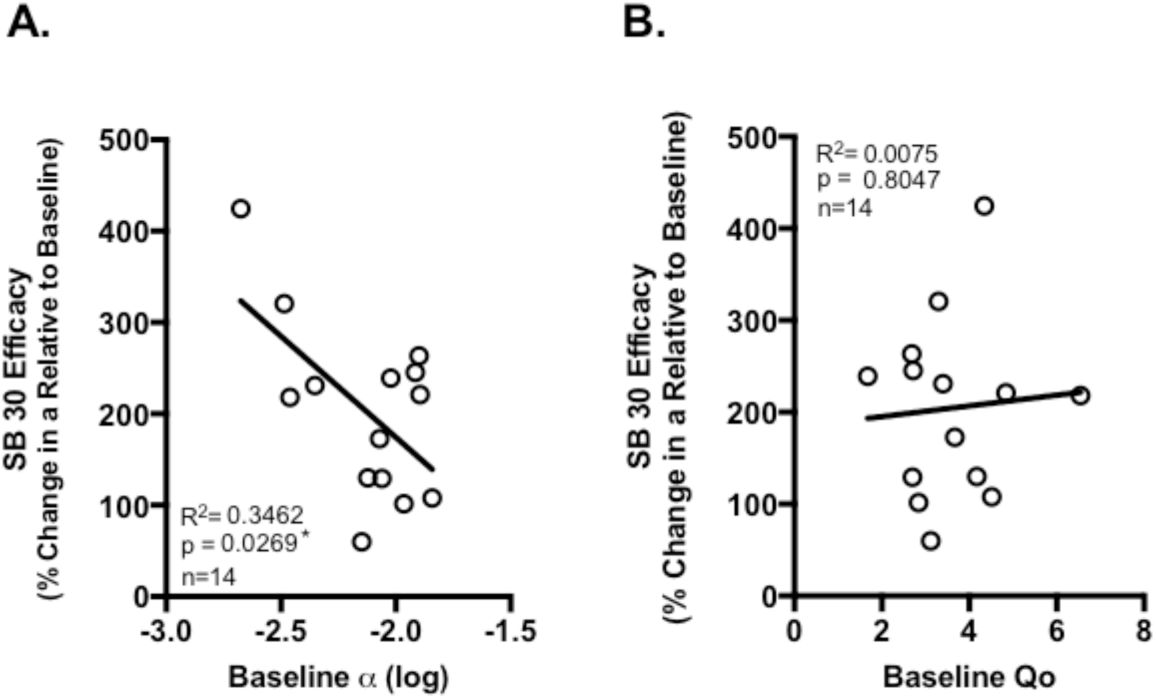
Baseline α values predict SB efficacy. A negative correlation was observed between baseline α values and SB (30mg/kg) efficacy indicating that SB was most effective in increasing demand elasticity in highly motivated rats (small α). B) Baseline Qo values did not predict the efficacy of SB in increasing demand elasticity.

### SB attenuates cue-elicited fentanyl seeking

To determine if OxR1 signaling mediates cue-elicited seeking of fentanyl, a subset of rats (n=10) underwent extinction training. Overall rates of responding decreased over the first 7 extinction sessions (Figure 4A; main effect of Session F_6, 54_ = 5.793, p=0.0001). Rats initially responded more on the active vs inactive lever (main effect of Lever Type F_1, 9_ = 66.5, p<0.0001). However, active vs inactive lever presses did not differ significantly by day 7 (although there was a trend; Time x Lever Type interaction F_6, 54_ = 4.165, p=0.0016; Bonferroni correction p<0.05). On averaged rats reached extinction criterion in 11 (±1.18) sessions.

**Figure 4.**
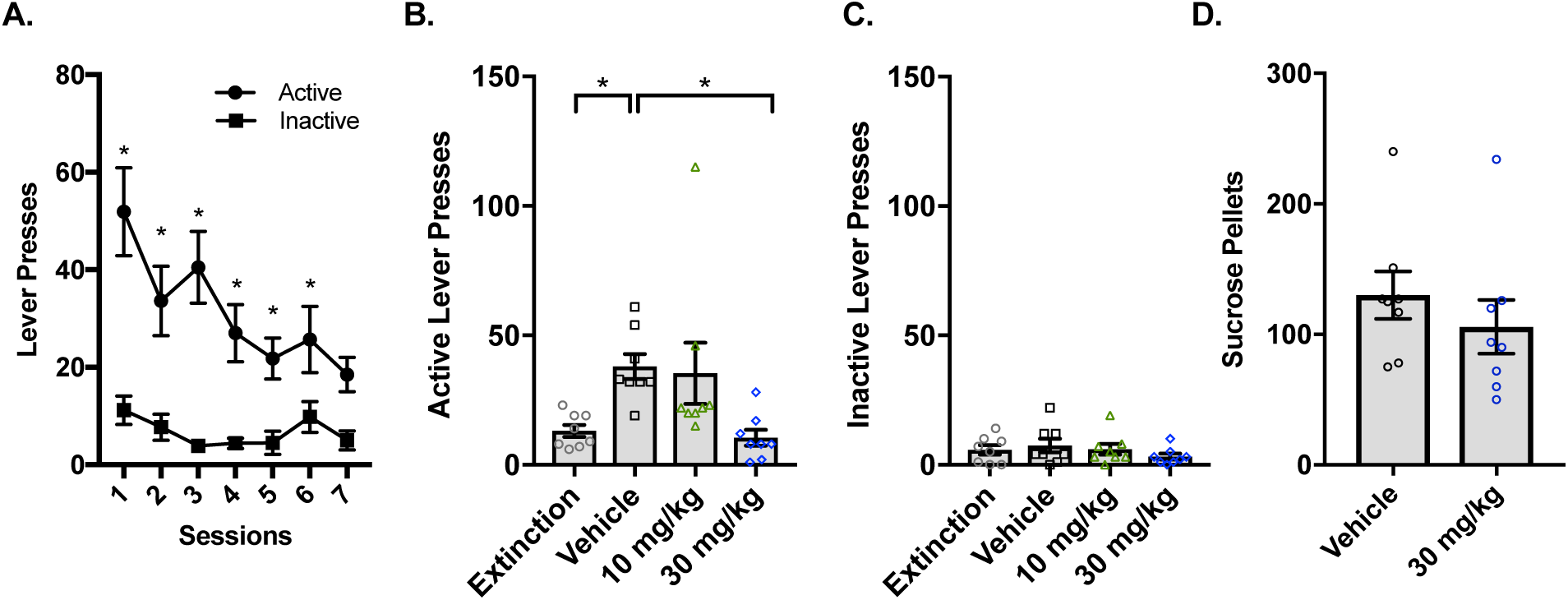
OxR1 antagonism attenuates cue-elicited drug seeking. Following BE testing, lever press responding was extinguished in a subset of rats. A) Active lever presses gradually decreased across the first 7 extinction sessions. B) The presentation of fentanyl-associated cues reinstated active lever pressing. SB at 30 mg/kg attenuated cue-induced reinstatement. C) SB had no effect on inactive lever presses. D) To confirm that the attenuation of cue-induced reinstatement by SB was not due to locomotor impairments, fentanyl experienced rats were trained to lever press for sucrose on an FR-1 schedule. Consumption of sucrose did not differ between SB (30 mg/kg) and vehicle treated rats. * p<0.05.

After reaching extinction criterion, rats (n=8) were given a systemic injection of vehicle or SB (10 or 30 mg/kg) prior to testing for cued reinstatement. SB pretreatment dose-dependently attenuated cue-induced reinstatement (Figure 4B; F_1.37, 9.61_= 6.796, p=0.0207). *Post-hoc* analysis revealed that the presentation of fentanyl-associated cues induced a significant reinstatement of drug seeking as measured by increased responding on the active lever (extinction vs vehicle; Dunnett's multiple comparisons test, p=0.0034). Compared to vehicle, SB (30 mg/kg) significantly reduced active lever presses (Dunnett's multiple comparisons test, p=0.0009). Changes in inactive lever responding were not observed (Figure 4C; F_3,21_=1.632, p=0.2121).

The effects of SB (30mg/kg) on sucrose self-administration were investigated to test for a potential confound of specificity and motor impairment (Reiner et al., 2018). Following their final cued reinstatement session, rats (n=8) were trained to self-administer sucrose pellets on an FR-1 schedule. Lever presses did not differ significantly between vehicle and SB (30 mg/kg) treated rats (Figure 4D; t_7_= 2.216 p=0.0623), indicating that lever pressing for a non-drug reinforcer was not impaired by SB treatment.

### Baseline α values predict cue-induced fentanyl seeking

Previous studies found that baseline α values predicted other DSM-V addiction-like behaviors, including cue-induced reinstatement of cocaine seeking (Bentzley et al., 2014; James et al., 2018a). Accordingly, we sought to determine if this were also true for fentanyl. Baseline α was negatively correlated with the degree of cue-induced fentanyl seeking (percent change in active lever presses from final extinction session) such that reinstatement behavior was greatest in highly motivated rats (Figure 5A; R^2^=0.4242, p=0.0414). In contrast, no correlation with reinstatement responding observed with baseline Qo (Figure 5B; R^2^=0.1522, p=0.2650).

**Figure 5.**
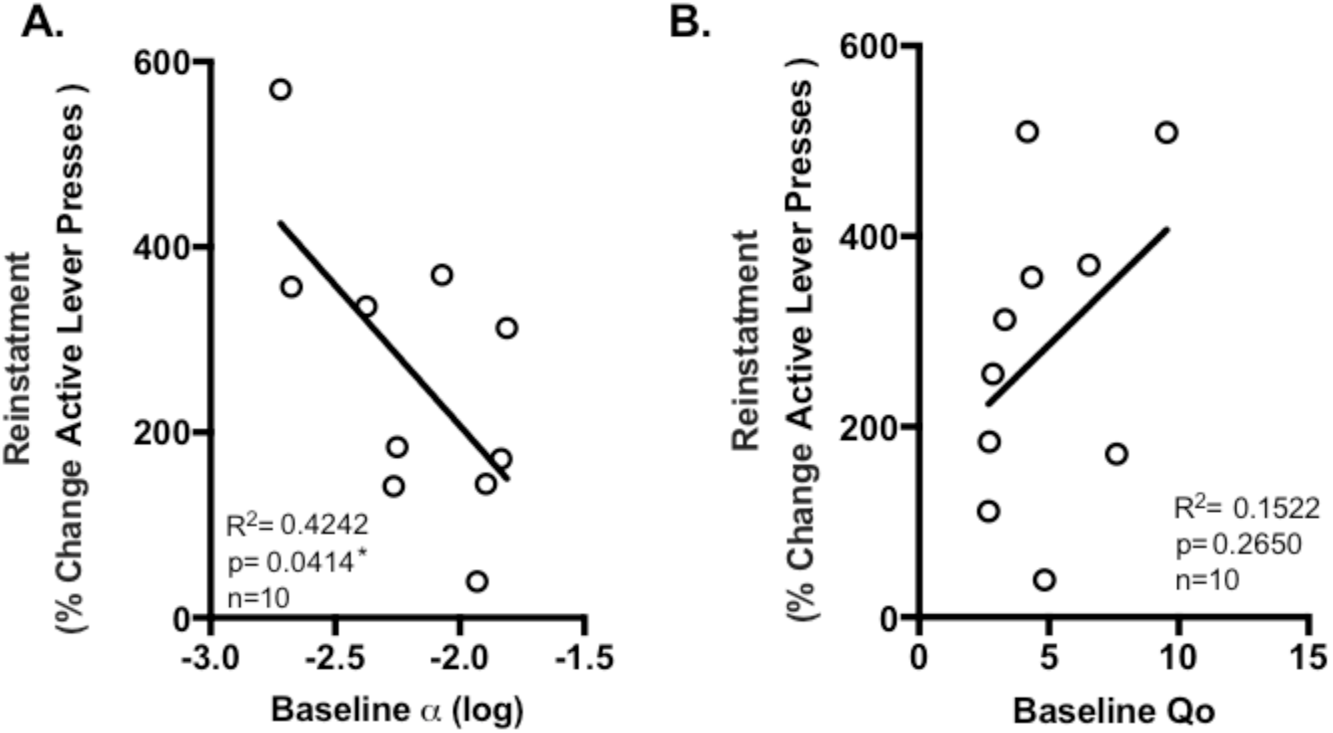
Baseline α values predicted cue-induced reinstatement of fentanyl seeking. The relationship between baseline α and Qo values and reinstatement (% change in active lever responses from final extinction session) was assessed. A) Baseline α was negatively correlated with reinstatement such that highly motivated rats (low α) showed greater drug seeking in response to cues. B) No relationship was observed between baseline Qo values and reinstatement responding.

## Discussion

The present study sought to determine the role of OxR1 signaling in demand for the opioid fentanyl using a novel within-session BE procedure. We found that systemic pretreatment with the OxR1 antagonist SB increased demand elasticity for fentanyl (decreased motivation) without affecting preferred consumption of fentanyl at null cost (Q_o_; hedonic setpoint). Baseline α values predicted SB efficacy, such that SB pretreatment was most effective at increasing demand elasticity in highly motivated rats. SB did not alter sucrose consumption in rats simultaneously receiving fentanyl indicating that increased demand elasticity following SB pretreatment is likely not due to sedation or motor impairment. In addition, SB dose dependently attenuated cue-induced drug seeking and baseline α values predicted the degree of reinstatement observed. These results implicate OxR1 signaling in fentanyl abuse and emphasize the utility of BE procedures in assessing addiction-related behaviors for this drug.

### A preferential role for OxR1 signaling in motivation

Here, we show that OxR1 blockade decreases motivation for fentanyl (increases α) without altering free consumption (Q_O_). These results align with previous reports showing a preferential role for OxR1 signaling in regulating consumption of drug and natural reward under high-effort schedules of reinforcement (Borgland et al., 2009; Espana et al., 2010; Hollander et al., 2012; Smith et al., 2009). For example, OxR1 blockade reduces cocaine consumption under progressive ratio (PR) but not FR-1 schedules of reinforcement (Boutrel et al., 2005; Espana et al., 2010; Smith et al., 2009). Consistent with these findings, SB increased demand elasticity (α) for cocaine without effecting preferred consumption at null cost (Qo) (Bentzley and Aston-Jones, 2015; James et al., 2018b). For natural reinforcers, such as sucrose and high fat diet, OxR1 blockade preferentially reduced consumption under high-effort schedules of reinforcement (Borgland et al., 2009; Cason and Aston-Jones, 2013). Together, these results support a role for OxR1 signaling in motivational verses hedonic processes.

There are, however, examples of OxR1 signaling regulating drug consumption under low-effort schedules of reinforcement. OxR1 receptor blockade reduces consumption of ethanol under low-effort schedules of reinforcement as well as homecage ethanol preference (Lawrence et al., 2006; Moorman and Aston-Jones, 2009; Moorman et al., 2017). However, these results were specific to ethanol-preferring rats (Lawrence et al., 2006; Moorman and Aston-Jones, 2009) and high-takers (Moorman et al., 2017). This may also explain why OxR1 blockade reduces consumption of heroin under both low and high schedules of reinforcement (Smith et al., 2009). This study did not separate rats into high and low responders making it unclear if the effect of SB on low-effort responding was driven by high takers. Using an identical within-session BE task, we previously found that systemic SB increased demand elasticity and reduced consumption of remifentanil at null cost (Porter-Stransky et al., 2017). In this study, α and Qo values were correlated making it difficult to determine whether effects on Qo were simply driven by changes in α. Nonetheless, these studies support a role for OxR1 signaling in drug demand and highlight the orexin system as an encouraging pharmacotherapy for drug addiction.

### OxR1 signaling and cue-induced reward seeking

The orexin system also seems to play a ubiquitous role in cue-induced reward seeking. Orexin neurons are activated in response to drug-associated cues and contexts (Dayas et al., 2008; Moorman et al., 2016; Plaza-Zabala et al., 2012). This is consistent with previously published reports showing that the OxR1 is necessary for cue-induced reinstatement of heroin, remifentanil, cocaine, ethanol, nicotine, and sucrose (Moorman et al., 2017; Plaza-Zabala et al., 2012; Porter-Stransky et al., 2017; Smith and Aston-Jones, 2012; Smith et al., 2009). Here, we show that OxR1 antagonism also attenuates cue-induced fentanyl seeking. The role for OxR1 signaling in motivation (above) is consistent with this role in relapse to drug seeking; evidence from our lab indicates that the OxR1 may play a particularly important role in the motivation afforded to drug-associated stimuli (Bentzley and Aston-Jones, 2015). Here, we show that OxR1 antagonism has no effect on high-rate responding for sucrose. This suggests the effects of SB on cue-induced reinstatement were not due to nonspecific drug effects. These results indicate that OxR1 signaling plays a strong role in cue-induced drug seeking and OxR1 signaling antagonist may be particularly useful in preventing relapse.

### Demand elasticity and the efficacy of orexin-based pharmacotherapies

We found here that baseline α values for fentanyl are negatively correlated with SB efficacy. This aligns with previous studies showing that OxR1 antagonism preferentially reduced drug seeking behaviors in highly motivated rats (James et al., 2018a; Moorman and Aston-Jones, 2009; Moorman et al., 2017). As previously stated, SB selectively reduces ethanol self-administration in high-ethanol responding rats (Moorman and Aston-Jones, 2009) and SB is most effective at decreasing cocaine motivation in rats with high baseline demand (James et al., 2018a). Highly motivated rats are also more sensitive to lower doses of SB (James et al., 2018b; Schmeichel et al., 2017). Together with the current study, these findings point to an increased reliance on OxR1 signaling for the maintenance of motivated behavior in highly motivated individuals. In addition, these results emphasize the utility of demand elasticity in identifying which individuals would most benefit from orexin-based pharmacotherapies.

### Demand elasticity and addiction propensity

Demand elasticity provides a powerful cross-species behavioral biomarker of addiction propensity (Bruner and Johnson, 2014; MacKillop et al., 2018; MacKillop et al., 2008; Pickover et al., 2016). Here, we show that baseline α values predict cue-induced fentanyl seeking. This is consistent with previously published reports showing that baseline α values for cocaine are predictive of other DSM-V relevant addiction-like behaviors including compulsive responding, drug seeking during initial absence, and cue-induced reinstatement (Bentzley et al., 2014; James et al., 2018a).

## Conclusions

Here, we show that OxR1 signaling regulates motivation to consume fentanyl and cue-induced fentanyl seeking. We also show that baseline α values predict SB efficacy and cue-induced fentanyl seeking. Taken together, our findings support the promise of OxR1 antagonist as a novel pharmacotherapy to combat the opioid epidemic. Furthermore, our results illustrate the usefulness of BE procedures in identifying individuals that may benefit best from orexin-based pharmacotherapies and those with the most severe cases of opioid abuse.

## Acknowledgements

We would like to thank Veronica Behman for her assistance in performing behavioral experiments for this study.

